# EEG Changes during Odor Perception and Discrimination

**DOI:** 10.1101/2022.12.12.520035

**Authors:** Marina Morozova, Alsu Bikbavova, Vladimir Bulanov, Mikhail A. Lebedev

**Affiliations:** V. Zelman Center for Neurobiology and Brain Restoration, Skolkovo Institute of Science and Technology, 121205 Moscow, Russia; VIBRAINT RUS LLC, 121205 Moscow, Russia; Faculty of Mechanics and Mathematics, Moscow State University, 119991 Moscow, Russia; Laboratory of Neurotechnology, I. M. Sechenov Institute of Evolutionary Physiology and Biochemistry, 194223 Saint-Petersburg, Russia

**Keywords:** olfaction, brain-computer interface (BCI), neurofeedback, electroencephalography (EEG), respiratory cycle

## Abstract

Brain-Computer Interfaces (BCIs) are devices designed for communication between the central nervous system and a computer. The communication can occur through different sensory modalities, and most commonly visual and auditory systems are used. We suggest that BCIs can be expanded by the incorporation of olfactory modality. To probe the modulation of EEG activity by odorants, we implemented two olfactory tasks: one that required attention to perception of odors, and the second one where participants discriminated consecutively presented odors. EEG recordings were conducted in 13 healthy participants while they performed the tasks following computer-generated verbal instructions. Respiratory activity was recorded to relate EEG modulations to the breath cycle. Theta activity responses were observed over the frontal EEG leads approximately 2 s after the inhalation. This theta activity was different depending on whether or not the inhaled air contained an odorant, and cognitive load also had an effect. We conclude that the frontal theta rhythm is reflective of the neural processing of odors. As such, it could be incorporated in the olfactory-based BCIs which take odors either as inputs or outputs. These BCIs could improve olfactory training needed for conditions like anosmia and hyposmia, and mild cognitive impairment.

## 1 Introduction

One of the clinical symptoms of COVID-19 is a sudden deterioration of olfaction, where COVID-19 has a greater impact on odor threshold detection than on odor identification (Le Bon et al., 2021). Electroencephalography (EEG) is potentially one of the tools for testing olfactory function in humans. Yet, in contrast to commonly used Mismatch Negativity response and P300 response, chemosensory event-related potentials (ERPs) are typically small, so it is challenging to analyze them. Schriever et al. (2017) analyzed the induced EEG-power changes during the interval 200–2000 ms after aroma onset in the frequency band 2 to 6 Hz, and found that these EEG patterns could be used to distinguish people with olfactory impairment from healthy individuals. The particular time-frequency parameters were chosen according to the previous studies (Huart et al., 2012) where the chemosensory event-related potentials (ERPs) to trigeminal and olfactory nerve stimulation were analyzed.

Although the respiratory cycle is critical for olfaction, no breath sensors were used in the majority of studies on olfaction. Only in a few studies the respiratory cycle was accounted for (Haehnel et al.,2011). Overall, there is still no clear understanding of the EEG changes related to the respiratory cycle, despite it being known that respiration affects functional brain connectivity (Perl et al., 2019).

Studies in animals have shown that theta oscillations occur during olfactory processing, and these oscillations are driven by the low-frequency burst firing of external tufted cells in the glomerulus (Kay et al., 2009). Additionally, odor-evoked gamma and beta oscillations are associated with odor learning, particularly in cognitive tasks (Beshel et al., 2007).

At the moment, the most common way to improve sensitivity to odors and restore olfaction is olfactory training (OT) (Addison et al., 2021). Moreover, a potentially neurofeedback could be a potentially useful tool to improve OT. In such an olfactory-based neurofeedback system, neural patterns occurring during processing of odors are decoded from brain recordings and presented to the user as visual feedback or feedback based on a different sensory modality. Yet, it is critical for implementing a neurofeedback system that neural responses to odors are well understood and properly decoded. It is still not well understood how olfactory-based neurofeedback could be derived from EEG recordings. In addition to neurofeedback, it is of interest how olfactory processing could be incorporated in the design of brain-computer interfaces (BCIs), the devices that sample brain signals, analyze them, and translate them into commands that are transmitted to external device, for example assistive devices for patients with sensory, motor and cognitive disabilities (Alonso-Valerdi et al., 2015). Most BCI implementations rely on user interaction with visual and/or auditory or visual stimuli. A few of implemented BCIs incorporate somatosensory signals, and none of them utilize olfactory or gustatory modalities. We propose that incorporating olfaction in BCIs should be explored as a potentially powerful way to modulate brain activity. Indeed, the olfactory system is a phylogenetically old network that connects to many brain areas. The olfactory system is unique neuroanatomically as it connects to the olfactory cortex without a thalamic relay.

The current study was motivated by three considerations relevant to the development of olfactory-based BCIs. First, we suggested BCIs with an olfactory output could be useful for certain applications, particularly the ones where relaxation and positive emotions are required. Second, we suggested that olfactory-based BCIs could be implemented using the P300 design where an olfactory stimulus triggers a cortical evoked response that could be further analyzed by the decoding algorithm. A variant of this idea is using visually evoked responses that match a particular odor. Third, we reasoned that the operation of different types of BCIs could be enhanced by adding odors that enrich the sensory stimuli, for example, adding odors to a virtual environment.

While these ideas appear plausible, better understanding of EEG patterns occurring during odor processing is needed. As a step toward attaining this understanding, we conducted a study where EEG was sampled in participants who discriminated odors.

## 2 Materials and methods

### 2.1 Experiment design

Thirteen healthy volunteers (6 males; age: 24.1 ± 5.8, mean ± SD) participated in the study. All subjects did not have a history of neurological diseases and reported having no significant changes in the sense of smell in the previous half-year period. All participants were confirmed to be normosmic using the Sniffin’ Sticks test 12 items (SST-12). All participants were native Russian speakers; during the experiment, the commands were given in Russian using computer audio. Experiments were approved by the local Ethics Committee of the Skolkovo Institute of Science and Technology, Moscow. All volunteers gave informed consent to participate in the study.

For the delivery of odors, we used Aroma Shooter® diffuser (Aromajoin Corporation, Japan) and six Aroma Cartridges including Caramel, Grass, Orange, Pine, Smoke, and Mint scents (identification numbers S-SW4, S-GN1, N-CT1, N-WD7, S-IM16, and N-HB21, respectively). We used the free Aroma Shooter® SDK implemented in the VIBRAINT Research software (VIBRAINT RUS LLC, Russia) which runs two olfactory exercises, namely, Random and Matrix.

During the first experimental condition, called Random, participants familiarized themselves with the aromas. The participants were given 100 trials where they responded to commands “Exhale” and “Inhale” after an odor was sprayed by the Aroma Shooter® diffuser using a 0.5 s long sprat. In half of the trials, no odor was delivered. The participants were instructed that no odor would be delivered in some trials. They were also instructed to memorize odors they felt because they would be required to discriminate them during the second part of the experiment. The participants were not given any additional information about the names of odors.

During the second experimental condition, called Matrix, participants were required to discriminate odors. The condition included 40-60 trials (median 50 trials), each consisting of the following sequence of steps: 1) an auditory command “New pair”; 2) commands “Exhale” and “Inhale” followed by a 0.5 s long spray of the first odorant from the Aroma Shooter® diffuser; 3) a 5 s period for the order to dissipate; 4) commands “Exhale” and “Inhale” followed by a 0.5 s spray of the second odorant from the Aroma Shooter® diffuser; 5) the command “Confirm match” after which the participant pressed “1” or “0” on the keyboard to report that the first and the second odorants were identical or different, respectively. The number of trials with the identical odorants in the pair was equal to the number of trials with different odorants.

### 2.2 Data acquisition and analysis

Throughout the experiment, the raw EEG data were collected using an NVX-36 amplifier (MKS, Russia). 22 EEG channels were recorded according to the international “10–20” system with sampling frequency 250 Hz using Ag/AgCl electrodes lubricated by an electrode gel. The ground electrode was attached to the FCz site. Two reference electrodes were placed on the left and right earlobes. The electrode impedance was kept below 15 kΩ. Respiration was measured with a nasal thermometric breath sensor TRSens (MKS, Russia). During the experiment, participants were asked to keep their eyes open. They sat in a comfortable position. The Aroma Shooter® diffuser was mounted 20 cm from the participant’s nose above.

EEG data were split into 4 s long epochs for processing. Each epoch started 1 s before the beginning of the inhale and ended 3 s after it. The beginning of the inhale was determined based on the measurements from the breath sensor. Each epoch was z-score standardized prior to the time-frequency analysis. Time-frequency analysis of the epochs was performed using the continuous Morlet wavelet transform (CWT) with the initial spread of the Gaussian wavelet set at 2.5/πω_0_ (ω_0_ being the central frequency of the wavelet). CWT was applied to a single epoch in the 1-40 Hz frequency band. To calculate event-related spectral perturbations (ERSPs), average time-frequency maps were calculated for the absolute values of single-trial records. Cluster-level statistical permutation test with one-way ANOVA as the test statistic was applied to compare time-frequency maps for odor and no-odor trials of the Random condition and the first and second odorants of the Matrix condition.

## 3 Results

### 3.1 Random condition

A cluster-level statistical permutation test was applied to compare CWT time-frequency maps of odor and no-odor trials of the Random condition. We found a significant increase in the frontal theta-range activity during the odor aroma trials as compared to the no-odor trials. This increase occurred during the interval from 1.5 to 2.5 s relative to inhale onset (fig. 1e). We calculated the mean power in the theta-range cluster for each epoch and found a statistically significant difference in the mean theta power for the comparison between the odor and no-odor trials (independent t-test statistic=-8.07, p-value=1.56e-15; fig. 1d) with a 10.5% increase in theta power. We also found a slight, yet statistically significant, correlation between the accuracy for the Matrix condition and the mean theta power in the Random condition (Pearson’s r=0.108, p-value=7.45e-05).

**Figure 1.**
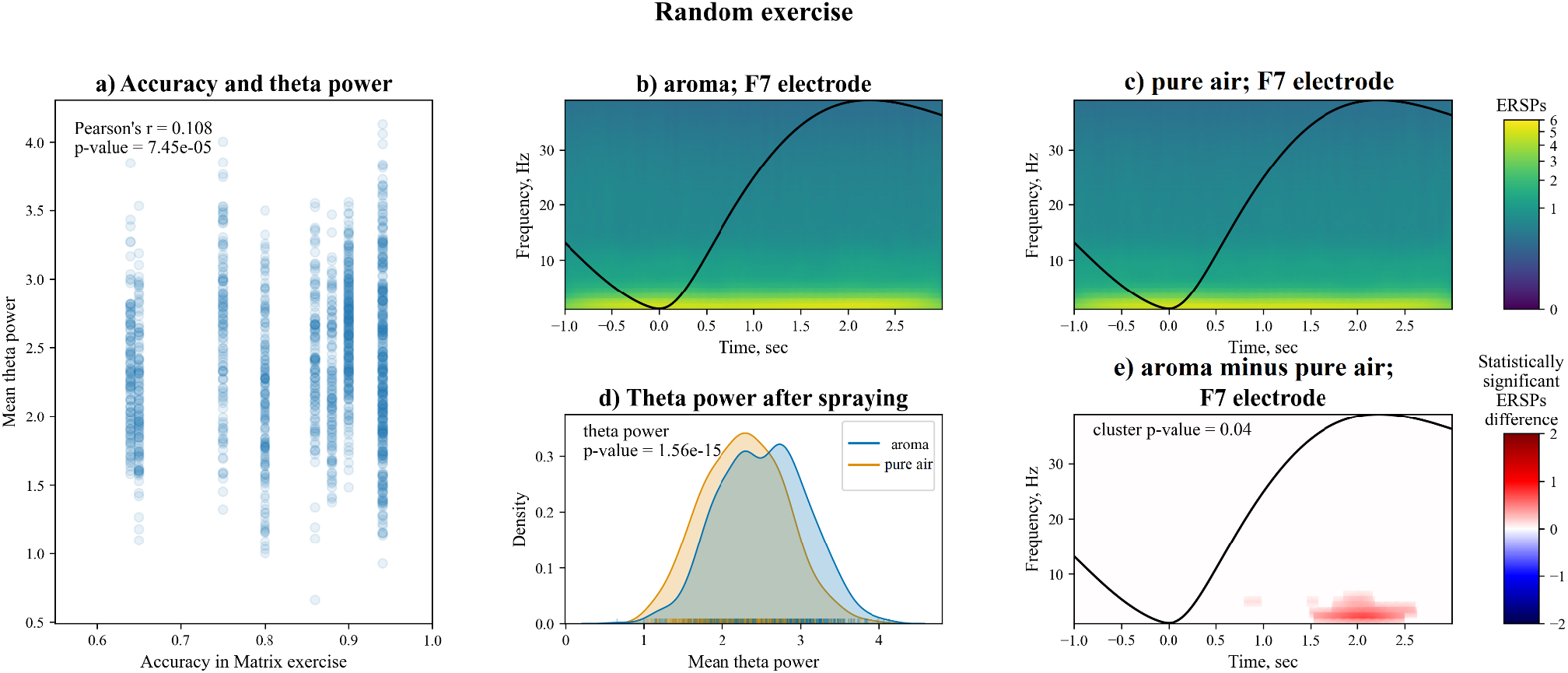
EEG analysis for the Random condition. A total of 1,300 trials were analyzed. In half of the trials, no odor was presented (no-odor trials), and the other half were odor trials. a) A scatterplot showing the correlation between the frontal theta power exhibited during the Random condition and report accuracy during the Matrix condition. b, c) The ERSPs for the odor (b) and no-odor (c) epochs. The black line represents the average breath cycle. d) Comparison of distributions of mean theta power for the odor and no-odor epochs. The distributions are calculated using kernel density estimation. e) The statistically significant time-frequency region of increased theta power during the odor trials compared to no-odor trials. The cluster was calculated using a cluster-level statistical permutation test for the absolute values of wavelets for the odor and no-odor trials. To measure the changes, ERSPs of the no-odor epochs (c) were subtracted from the ERSPs of the odor epochs (b). The black line represents the average breath cycle.

### 3.2 Matrix condition

Here we compared CWT time-frequency maps for the first and second odors epochs of the Matrix condition. A cluster-level statistical permutation test was used. We found a significant increase in the frontal theta activity. Besides the changes in the theta activity, we found a slight decrease in beta power during the second odor (fig. 2e). The calculated mean power in the theta-range cluster was statistically significantly higher during the second odor (independent t-test statistic=-3.4, p-value=0.00055; fig. 2d) with a 4.5% increase on average. A small but statistically significant correlation was observed between the accuracy in the Matrix condition and the mean theta power in the same condition (Pearson’s r=0.1, p-value=0.00057).

**Figure 2.**
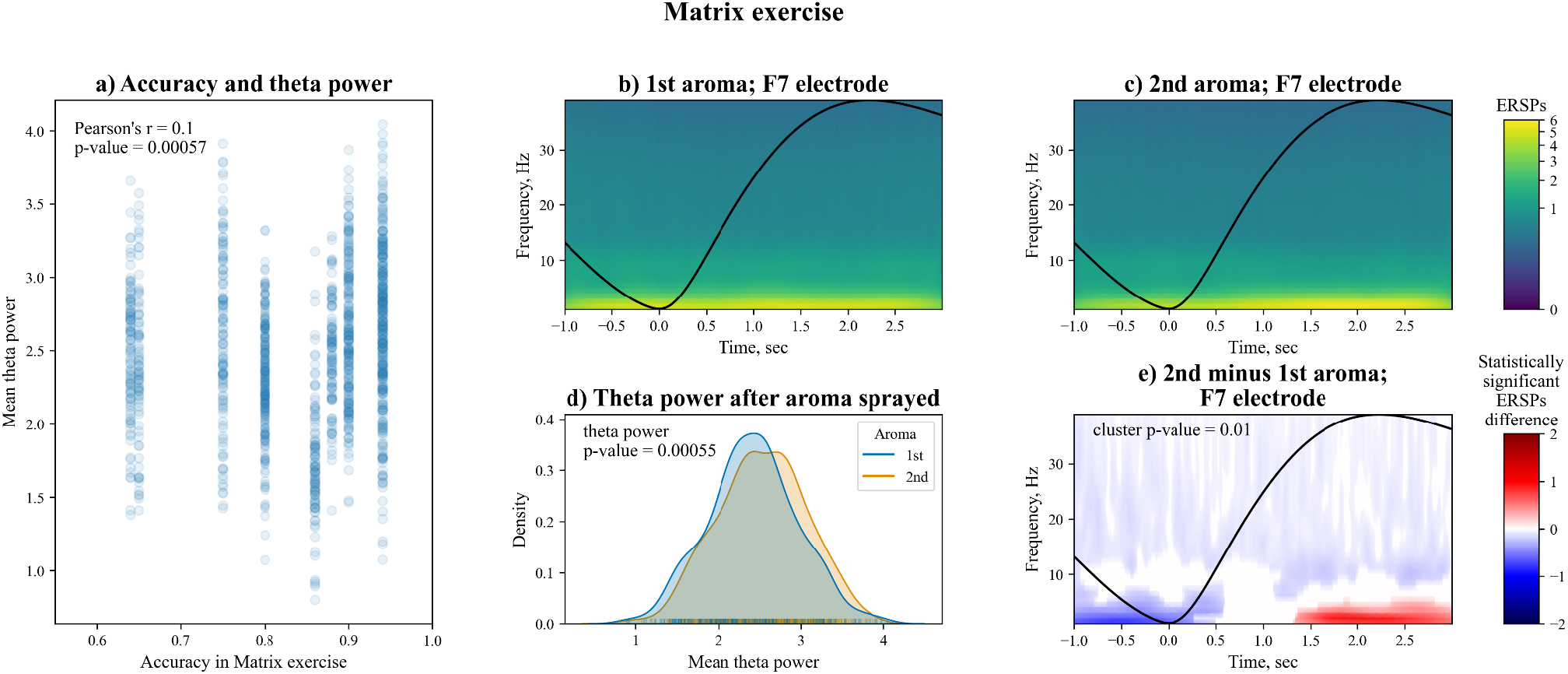
Analysis of EEG changes during the Matrix condition. A total of 1,300 trials were analyzed. In this analysis, epoch pairs were considered, which corresponded to the presentation of first and second odorants. a) The scatterplot showing the correlation between the frontal theta power and report accuracy. b, c) The ERSPs for the first (b) and second (c) aroma odors. The black line represents the average breath cycle. d) A comparison of the distributions of mean theta power during the first and second odors. The distributions were calculated via kernel density estimation. e) The period where a statistically significant increased theta power was found. The clusters were calculated using a cluster-level statistical permutation test between the absolute values of the wavelets for first and second odors. To measure the change, ERSPs for the second (c) odor were subtracted from the ERSPs for the first (b) odor. The black line represents the average breath cycle.

## 4 Discussion

We found a significant increase in the frontal theta power during the period when healthy participants inhaled odorants as compared to inhaling air without any odorants (fig. 1). The participants were instructed to pay attention to their olfactory perceptions, but they were not required to name or discriminate the perceived odors.

We also found a significant increase in the theta power when the participants compared two consecutively presented odorants and reported whether or not they were different (fig. 2).

The correlations of the theta power with, firstly, the presence/absence of aroma and, secondly, cognitive load provide the opportunity to use the theta power as an objective EEG-based metric for a machine learning model used in the neural interfaces. The model can be adjusted to give an objective score to estimate the threshold of perception of a smell and the level of focus to an olfactory task.

These results could be extended to building an olfactory-based BCI where frontal theta-power changes related to perceived odors are converted into an output signal (e.g. visual feedback). Conversely, frontal theta-power could be converted into an olfactory neurofeedback where odor type and intensity represent changes in the EEG. Such BCIs could be useful for rehabilitation of people with olfactory disabilities. Additionally, rehabilitation of age-related mild cognitive impairment is also a potential field of application.

## 6 Author Contributions

VB contributed to constructing the experimental setup and writing the software. Other authors equally contributed to the development of the experiment design, data collection and analysis, and writing of the manuscript.

## 8 Data Availability Statement

The dataset generated and analyzed for this study is available upon request.

